# Larval ecology and bionomics of *Anopheles funestus* in highland and lowland sites in western Kenya

**DOI:** 10.1101/2021.08.04.455104

**Authors:** Isaiah Debrah, Yaw A. Afrane, Linda Amoah, Kevin O. Ochwedo, Wolfgang R. Mukabana, Daibin Zhong, Zhou Guofa, Ming-Chieh Lee, Shirley Onyango, Edwin O. Magomere, Harrysone Atieli, Andrew K Githeko, Guiyun Yan

## Abstract

**Background:** *An. funestus* is a major Afrotropical vector of human malaria. This study sought to investigate the larval ecology, sporozoite infection rates and blood meal sources of *An. funestus* in western Kenya.

**Methods:** Larval surveys were carried out in Bungoma (Highland) and Kombewa (lowland) of western Kenya. Aquatic habitats were identified, characterized, georeferenced and carefully examined for mosquito larvae and predators. Indoor resting mosquitoes were sampled using pyrethrum spray catches. Adults and larvae were morphologically and molecularly identified to species. Sporozoite infections and blood meal sources were detected using real-time PCR and ELISA respectively.

**Results:** Of the 151 aquatic habitats assessed, 62/80 (78%) in Bungoma and 58/71(82%) in Kombewa were positive for mosquito larvae. Of the 3,193 larvae sampled, *An. funestus* larvae constitute 38% (1224/3193). Bungoma recorded a higher number of *An. funestus* larvae (85%, 95%, CI, 8.722-17.15) than Kombewa (15%, 95%, CI, 1.33-3.91). Molecular identification of larvae showed that 89% (n=80) were *An. funestus*. Approximately 59%, 35% and 5% of *An. funestus* larvae co-existed with *An. gambiae s*.*l, Culex spp* and *An. coustani* in the same habitats respectively. Of 1,221 *An. funestus s*.*l* adults sampled, molecular identifications revealed that *An. funestus* constituted 87% (n=201) and 88% (n=179) in Bungoma and Kombewa, respectively. The *Plasmodium falciparum* sporozoite rate of *An. funestus* in Bungoma and Kombewa was 2% (3/174) and 1% (2/157), respectively, and the human blood index of *An. funestus* was 84% (48/57) and 89% (39/44) and for Bungoma and Kombewa, respectively.

**Conclusion:** Man-made ponds had the highest abundance of *An. funestus* larvae. Multiple regression and principal component analyses identified the distance to the nearest house as the key environmental factor associated with the abundance of *An. funestus* larvae in aquatic habitats. This study serves as a guide for the control of *An. funestus* and other mosquito species to complement existing vector control strategies.

## Introduction

Malaria is still the most devastating vector-borne disease in sub-Saharan Africa, contributing approximately 215 million cases in 2019, which accounted for about 94% of all global cases [1]. Anti-vectorial programmes consisting mainly of long-lasting insecticide-treated nets and indoor residual spraying have contributed immensely towards reducing malaria incidence and mortality in malaria-,endemic areas of sub-Saharan Africa, which are characterized with having high entomological inoculation rates (EIR, infective bites per person per year) [1–3]. Kenya is noted among 17 countries estimated to have attained a reduction in malaria incidence in 2020 compared to 2015 [1]. Despite this unprecedented success in the fight against malaria, the Global Technical Strategy (GTS) 2020 milestone for reducing mortality and morbidity has not been achieved globally [1].

Currently, there is no “silver bullet” to successfully achieve the elimination and eradication goal outlined by the GTS. An important component of malaria vector control that needs reconsideration in the malaria elimination and eradication agenda is larval source management. Historical records have shown that a key component of malaria eradication efforts in Israel, Italy, and the United States of America was source reduction through larval habitat modifications [4]. In malaria-endemic areas of Africa, the use of insecticide-treated net, when combined with larval control, has been predicted to reduce the number of adult mosquito emergence by 50% and subsequently decrease the entomological inoculation rate (EIR) up to 15 to 25-fold [5]. However, the use of larval control strategies requires adequate knowledge of the larval ecology of the vectors, as well as better characterization of their breeding habitats in different ecological settings [6]. Targeting the most productive breeding habitats for larval control can be cost-effective, depending on the anopheline species, heterogeneity of aquatic habitats, and proximity to human dwellings [7,8].

*Anopheles funestus sensu stricto* (s.s) (hereafter, *An. funestus*) is a major Afrotropical vector of human malaria, exhibiting anthropophilic, and endophilic behaviours [9,10]. In western Kenya, *An. funestus* is one of the principal vectors of human malaria, owing to its high resistance to pyrethroids used for bed net impregnation, high sporozoite rate, and persistence in indoor malaria transmission [11,12]. Historically, western Kenya witnessed a decline in *An. funestus* populations after the introduction of insecticide-based control tools [13] and this reduction was observed for some time until a resurgence was reported a decade ago [12,14]. While this resurgence might be influencing malaria transmission in western Kenya, very few studies have examined and characterized the larval habitats of *An. funestus*. Unlike reports on *An. gambiae* s.s. and *An. arabiensis* in this region, few studies have found *An. funestus* larvae in large permanent habitats with thick aquatic vegetation and algae [15,16].

*Anopheles* mosquitoes breed in a range of aquatic habitats and assessing findable habitats is key in controlling immature stages of vectors. However, locating breeding habitats of *An. funestus* is difficult, as it infrequently breeds in notable habitats with other malaria vectors. Hence, a better understanding and characterization of the breeding habitats and ovipositional behavioural patterns of *An. funestus* in endemic areas is crucial before larval source management can make malaria elimination and eradication feasible. In this study, we characterized the aquatic habitats, larvae abundance, and examined the sporozoite infection rates and origin of blood meals of *An. funestus* in highland and lowland areas of western Kenya.

## Materials and Methods

### Study sites

This study was conducted in two distinct locations, a highland town (Bungoma) and a lowland town (Kombewa), situated about 55 km apart in western Kenya (Fig 1).

**Fig 1:**
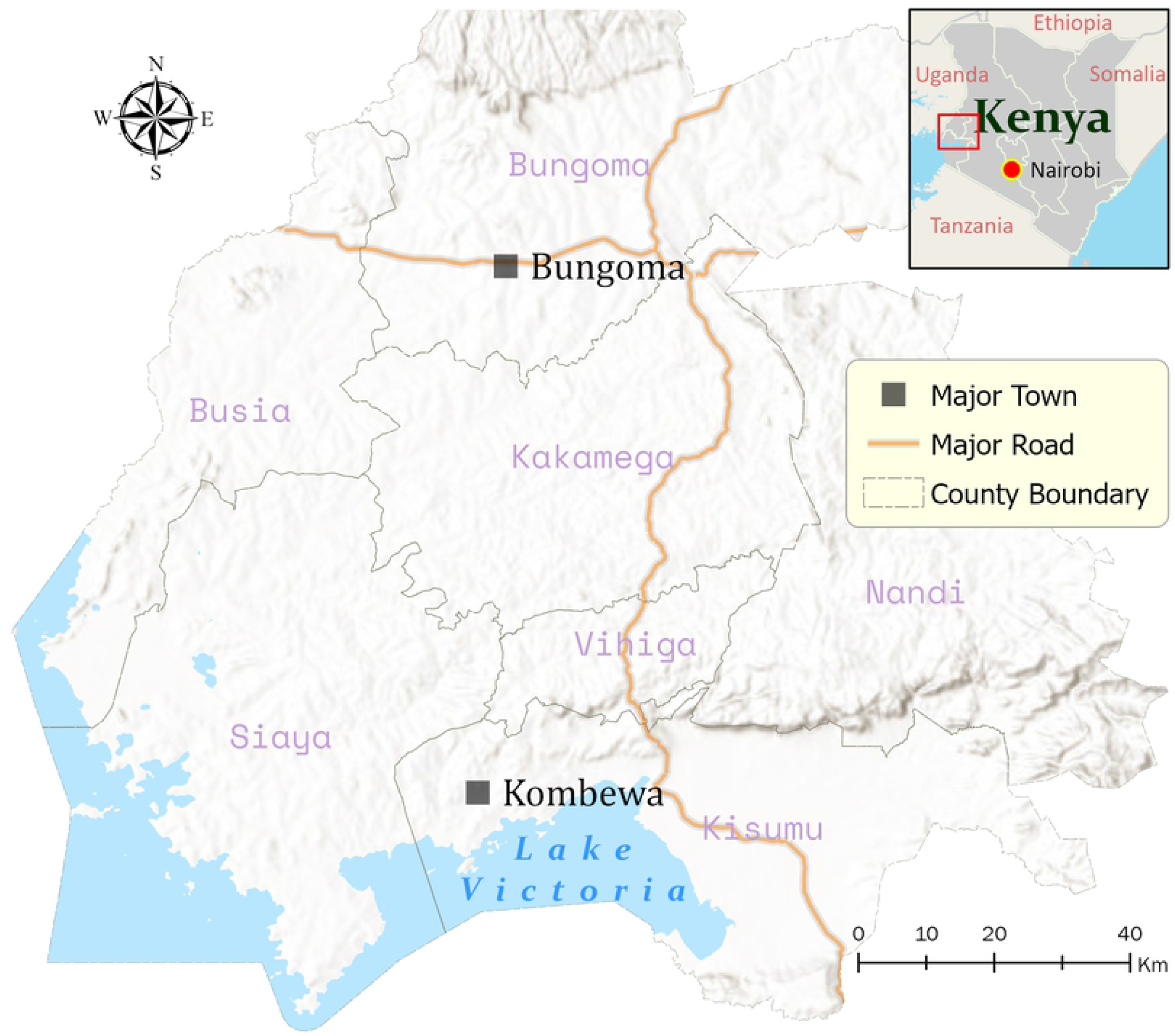
Map of the study areas in western Kenya.

Bungoma [00.54057°N, 034.56410°E, 1386–1,545m above sea level (asl)] is characterized by a perennial malaria outbreak. The mean annual rainfall and temperatures are 150 mm and 22.5 °C respectively [17]. The area previously had *Plasmodium falciparum* malaria episodes > 45% with hospitalization rate up to 55% [18]. Agricultural production of crops (tobacco, cereals, sugar cane, onions and other vegetables) and raising of farm animals (cattle, sheep and goat) and poultry form the backbone of the rural economy of Bungoma. The principal vectors of human malaria in Bungoma are *An. funestus, An. gambiae* and *An. arabiensis* [19,20].

Kombewa (34^0^ 30’E, 0^0^ 07’N, 1150 -1300 m asl) is located in Kisumu County, a lowland area with slow water drainage, in the Lake Victoria basin. This area experiences two rainy seasons: a long season from March to May characterized by peak malaria transmission and a short season between October and November. The mean annual rainfall and temperatures are 1200 mm-1300 mm and 20°C -35°C respectively [21]. There is, however, yearly variation in the rainfall pattern in the region. It is a hyperendemic malaria zone with a *P. falciparum* parasitemia rate of 57.5% [12] and EIR of 31.1 infective bites per person per year [11]. *An. funestus* has been reported to be the principal malaria vector predominating in this study area [11,22].

### Larval sampling

Larval surveys were carried out monthly from November 2019 to November 2020. Water bodies were identified within 2km of the study villages. Larvae were sampled using standard dippers (350 ml) and 10L bucket [23]. A maximum of 20 dips was taken at each habitat to identify the presence or absence of larvae and aquatic predators. Mosquito larvae were sampled along the edges of the habitats after waiting for 3-5 minutes for larvae to rise to the water surface if there were any. *An. funestus sensu lato (s*.*l)* larvae were preserved in absolute ethanol for subsequent identification using molecular analysis. Larvae were morphologically identified following referenced keys [24,25]. All water bodies were georeferenced using a global positioning system installed in ODK software in Android Samsung tablet (Version SAM-T380). Larval habitats variables were characterized following a questionnaire that was imputed using ODK software.

### Characterization of aquatic habitats

All potential water bodies for mosquito larvae were identified and classified as: man-made pond, natural pond/rain pool, drainage ditch, swamp/marshes and tire track/road paddle. Man-made ponds/pits were made purposely for moulding clay pots in Bungoma, whereas in Kombewa, they were dug for sand winning. Aquatic habitats were classified as permanent if they could hold water for more than three weeks. Environmental variables including the presence of vegetation, category of vegetation, the height of vegetation, vegetation coverage, substrate type, and distance to a nearest house, habitat dimensions (length, width and depth), water physics (flow status and clarity) and present or absent of aquatic predators were recorded for each water body. In addition, the surrounding land use type (cultivated land/cropland, grassland/pasture, wetland/swamp and road) were also recorded.

The vegetation cover was classified as: emergent, freely floating, submerged, and no vegetation. The height of vegetation coverage was measured using a meter stick and grouped into < 5, 6-10, and > 10 m. The distance to the nearest dwellings was estimated and grouped as (A) < 100 m, (B) 100-200 m, and (C) 201-500 m. The types of plants and predators in the habitats identified using picture charts. The dimensions (length, width and depth) of each habitat were measured using a meter stick, and the average depth was recorded. An Aquafluor^™^ meter (Model: 8000-010, Turner Design, San Jose, CA, USA) was used to measure cyanobacteria (blue-green algae) levels in water samples from the aquatic habitats.

The clarity of the water was observed and classified as (A) clear, (B) opaque, (C) cloudy, and (D) muddy/brownish. The substrate type was classified as (A) mud/dirty, (B) sand and (C) stone. Land-use types for the surrounding habitats were described as (A) cultivated land/cropland (farmland use for cultivation of food crops and other crops), (B) grassland/pasture (lands with suitable grasses for grazing animals) and (C) wetland/swamp (surroundings characterized by mostly aquatic plants species covered by water/ low-lying ground not cultivated and covered with water) and (D) road (land meant for passage of vehicles and people).

### Adult sampling

To find out whether larval abundance correlated with the adult *Anopheles* density, adult mosquitoes were collected indoors using the pyrethrum spray catches [26] from randomly selected houses near the aquatic habitats from November 2019 to August 2020. Collections were carried out in the morning hours between 06:30 to 10:00 h. The physiological status of the anophelines was classified into blood-fed, unfed, half gravid and gravid. Samples were stored in 1.5 ml Eppendorf tubes containing silica gel desiccant and cotton wool at -20° C in the Sub-Saharan Africa International Centre of Excellence for Malaria Research, Tom Mboya University College, Homa Bay, Kenya, for further analysis.

### DNA extraction and molecular identification of species

Adult mosquitoes were cut into two parts to separate the head and thorax from the abdomen. DNA was extracted from the head and the thorax using the Chelex^®^-100 method [27], while the abdomen was preserved for blood meal analysis. DNA was extracted from the larvae of *An. funestus s*.*l*. using the ethanol precipitation method [28]. Molecular identification of sibling species of the *An. funestus* group was performed using multiplex polymerase chain reaction (PCR) by amplifying the polymorphic ITS2 region of ribosomal DNA using species-specific primers for *An. funestus, An. rivulorum, An. vaneedeni, An. parensis, An. leesoni, An. rivulorum-like* by following already developed protocols [29,30]. Of the *An. funestus s*.*l* specimens that failed to amplify by PCR after three attempts, 65 were randomly selected and sent to the University of California, Irvine, for sequencing using the Sanger sequencing method. The ITS2 region of nuclear ribosomal DNA was amplified using the forward primer ITS2A (TGTGAACTGCAGGACACAT) and the reverse primer ITS2B (TATGCTTAAATTCAGGGGGT); amplicons were sequenced using ABI Big Dye Terminator Cycle Sequencing Kits, as described by Daibin et al., [31].

### *Plasmodia* species genotyping using multiplex RT-PCR

A TaqMan assay was used to detect *Plasmodia* species (*P. falciparum, P. ovale* and *P. malarae*) infections in the DNA samples as previously described [32,33].

### Blood meal analysis

Direct enzyme-linked immunosorbent assays (ELISA) was used to detect the origin of blood meal in the abdomen of blood-fed *Anopheles* mosquitoes following existing protocol [34].

### Ethical Statement

This study was approved by Maseno University’s Ethics Review Committee (MUERC/00778/19). Verbal consent was sought and granted from heads of households and owners of farmlands where the adult mosquitoes and larvae were sampled.

### Data analysis

Data were analyzed using SPSS software (Version 21 for Windows, SPSS Inc., Chicago, IL) and Graph Pad Prism V.8.0.1. The relative abundance of each species was expressed as a percentage of the number of larvae per species divided by the total number of larvae collected for all species combined per habitat type. The relative abundance of *An. funestus* was calculated as the number of *An. funestus* larvae from a specific habitat divided by the total number of *An. funestus* larvae in all samples from a habitat type.

The types of breeding sites, the number of mosquito larvae sampled and species, and the number of aquatic predators were presented in tables and figures. The Kruskal–Wallis test was used to compare the number of larvae in different habitat types and for samples having more than two groups: habitat type (man-made pond, natural pond/rain pool, drainage ditch, swamp/marshes, tire/road paddle), distance to the nearest house (<100, 100-200, 201-500), water clarity (clear, opaque, brownish/muddy, cloudy), aquatic plant species in a habitat [*Pennisetum purpureum* (elephant grass), *Schoenoplectus californicus*, others/unknown], and land-use type (cultivated land/cropland, grassland/pasture, wetland/swamp, road). The Mann–Whitney U test was used to compare samples with two variables: vegetation (present or absent), category of vegetation (emergent or non-emergent), water flow status (stagnant or flowing water), and predators (present or absent). Multiple regression and principal components analyses were used to identify environmental variables associated with the abundance of *An. funestus* in the aquatic habitats. The human blood index (HBI) was calculated as the percentage of *Anopheles* mosquitoes that fed on humans over the total number of blood-fed *Anopheles* for which the blood meal origins were determined. The sporozoite rate of *P. falciparum* was calculated as the proportion of *Anopheles* tested for sporozoites over the total genotyped. Spearman’s correlation (*r*_*s*_) was used to find the relationship between the adult *An. funestus* population and the larval density at each study site. The results were considered statistically significant at *P* <0.05. For the sequence data analysis of the *An. funestus s*.*l*. specimen that failed to amplify by PCR, the de novo assembly of reads was performed using geneious software [35]. Basic Local Alignment Search Tool (BLAST) was used to identify sequence similarities against sequences in GeneBank. Sequences with high identity score or low E-value were retrieved and used in the construction of a phylogenetic tree to identify the unknown. Evolutionary analyses were conducted in MEGA X after basic alignment using ClustalW algorithm [36]. All sequences of ITS2 are available at GenBank under accession numbers MZ435355-MZ435414.

## Results

### Distribution of *An. funestus* larvae and other mosquito larvae in Bungoma and Kombewa

A total of 151 mosquito aquatic habitats were assessed. Of these, 62/80 (78%; 95% CI: 0.68-0.87) and 58/71 (82%; 95% CI: 0.73-0.91) in Bungoma and Kombewa were positive for mosquito larvae, respectively. The number of aquatic habitats with *An. funestus* larvae in Bungoma was 55/80 (69%; 95% CI; 0.58-0.79), whereas 23/71 (32%; 95% CI: 0.21-0.43) were in Kombewa. In all, *An. funestus* larval habitats constituted 65% (n=78) of the mosquito-positive habitats, whereas *An. gambiae s*.*l, An. coustani*, and *Culex spp* positive habitats made up of 57% (n=69), 14% (n=17) and 36% (n=43), respectively.

A total of 3,193 mosquito larvae (39% *An. funestus*, 30% *An. gambiae* s.l., 28% *Culex* spp. and 3% *An. coustani*) were collected from the various habitats in Bungoma and Kombewa. Bungoma had a higher number of *An. funestus* larvae (85% 95%, CI, 8.722-17.15) than Kombewa (15% 95%, CI, 1.33-3.91). Approximately 59% of *An. funestus* larvae were collected from breeding sources with co-existing *An. gambiae s*.*l* larvae in the same habitats (Fig.2). Similarly, 35% and 5% of *An. funestus* larvae were found co-existing with *Culex spp*., and *An. coustani* larvae, respectively (Fig.2). No pupae were identified during the sampling. Molecular results on the sibling species of the *An. funestus* group larvae revealed that 89% (n=80) were *An. funestus*, 11% (n=9) were *An. rivulorum* and 1% (n=1) was *An. sp*.*9*.

**Fig 2:**
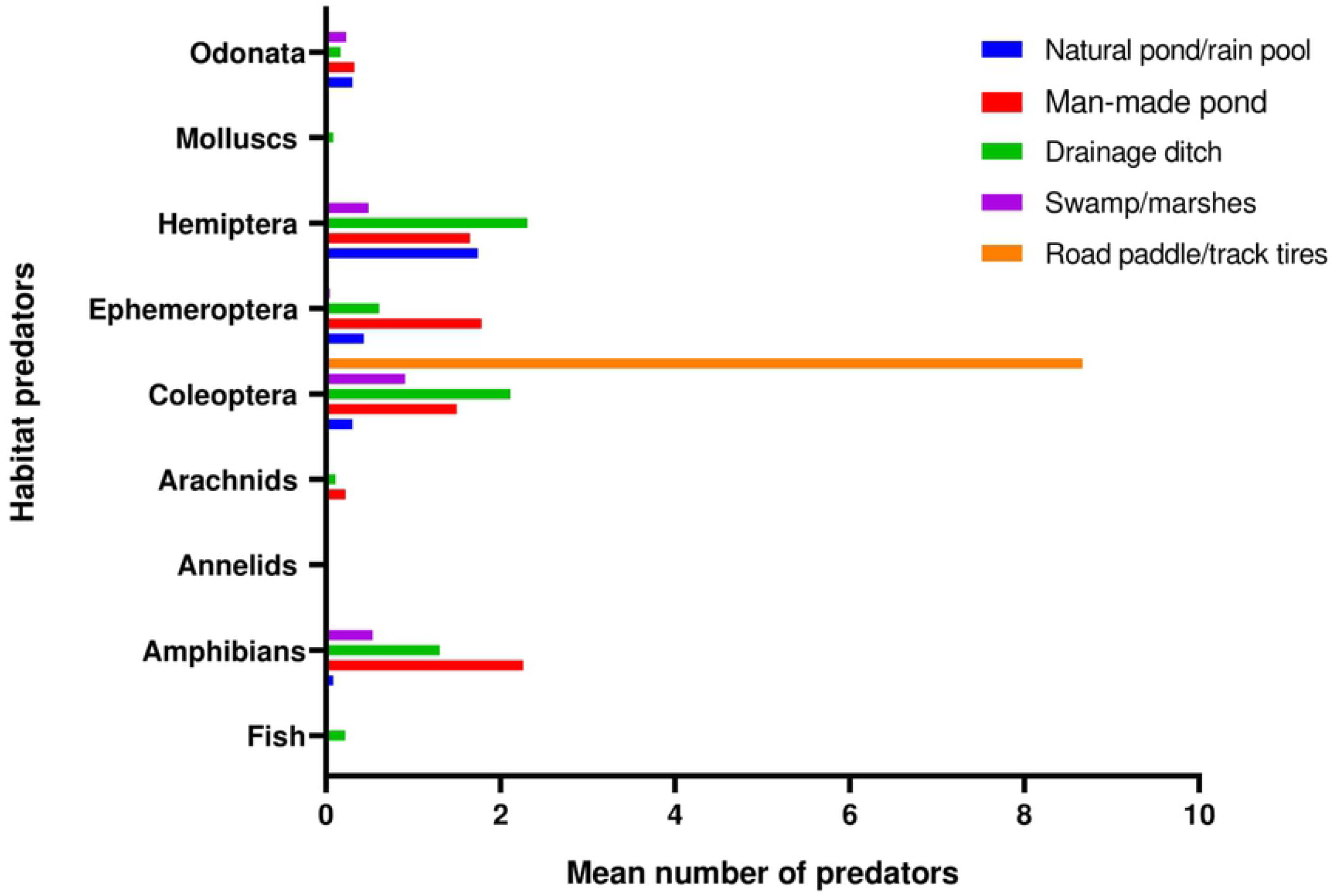
Proportion of *An. funestus* larvae shared with other mosquitoes in the larval habitats.

### Characteristics of mosquito larval habitats

*An. funestus* larvae were found in various habitats, with or without vegetation: man-made ponds, natural ponds/rain pools, drainage ditches, swamp/marshes, and tire/road paddles. There were no significant differences in *An. funestus* larval density between the various habitat types of *An. funestus* (χ2= 8.917, df=4, *P*=0.063). However, man-made ponds comprised the highest number of *An. funestus* positive habitats (36%, n=28) and had the highest proportion of larvae with *An. funestus* in Bungoma (53%, n=553) and in Kombewa (61%, n=115) (Table 1). Field observations showed that man-made ponds constituted the main permanent aquatic habitat type in the study areas. This was followed by swamp/marshes, natural ponds/rain pools, drainage ditches and tire/road paddles, at frequencies of 27%, 19%, 15%, and 3%, respectively. There was no significant difference in the means among the various predators found in the larval habitats (*P*=0.05). Fig 3 shows the mean distribution of the various predators.

**Table1:** Distribution of mosquito larvae collected from aquatic habitats in Bongoma and Kombewa, November 2019 to November 2020

| Study site | Habitats | No. of habitats | Larval Counts per Mosquito Species |  |  |  |
| --- | --- | --- | --- | --- | --- | --- |
|  |  |  | *An. funestus*<br>s.l n (%) | *An. gambiae*<br>s.l n (%) | *An. coustani*<br>i n (%) | *Culex*<br>spp n (%) |
| Bungoma | man-made pond | 26 | 553 (53) | 81(37) | 0(0) | 15(47) |
|  | natural ponds/rain pools | 12 | 100 (10) | 2(1) | 0(0) | 0(0) |
|  | drainage ditches | 19 | 374(36) | 120(54) | 0(0) | 3(9) |
|  | swamp/marshes | 21 | 8(1) | 18(8) | 0(0) | 14(44) |
|  | tire/road paddle | 2 | 0 | 0(0) | 0(0) | 0(0) |
|  | Total | 80 | 1035 | 221 | 0 | 32 |
| Kombewa | man-made ponds | 20 | 115(61) | 67(9) | 33(32) | 162(19) |
|  | natural ponds/rain pools | 11 | 25(13) | 249(34) | 24(23) | 188(21) |
|  | drainage ditches | 17 | 40(21) | 286(39) | 47(45) | 489(56) |
|  | swamp/marshes | 22 | 9(5) | 137(18) | 0(0) | 34(4) |
|  | tire/road paddle | 1 | 0(0) | 0(0) | 0(0) | 0(0) |
|  | **Total** | **71** | **189** | **739** | **104** | **873** |

**Fig 3:**
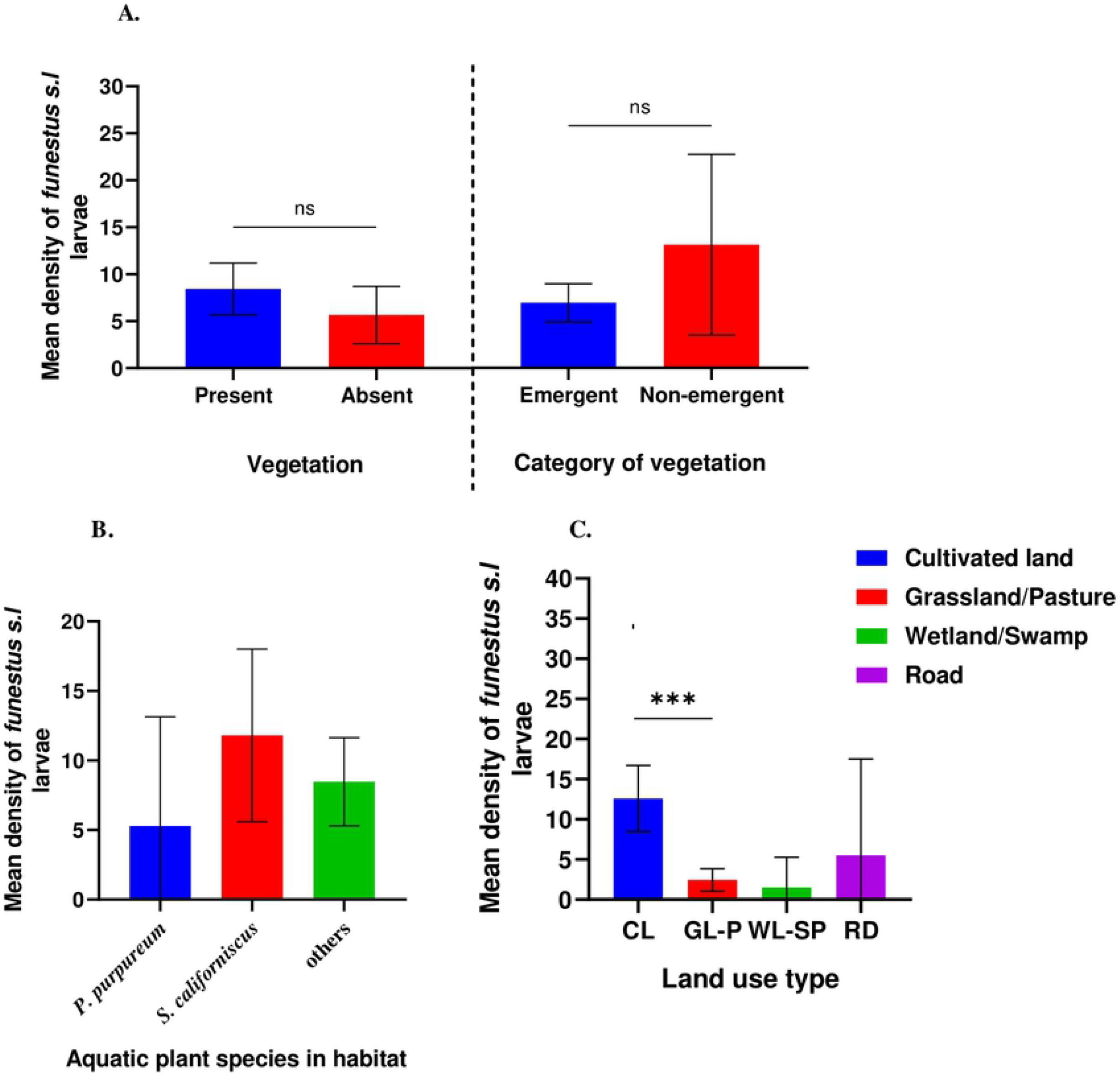
Distribution of aquatic predators in the various habitats. Odonata: damselfly/dragonfly, Molluscs: snails/slugs/mussels, Hemiptera: backswimmer/giant water bugs/cacidas, Ephemeroptera: mayfly, Coleoptera: water beetles/weevils, Arachnids: spiders/ticks/mites, Annelids: Segmented worms, Amphibians: frogs/toads/tadpoles, Fish: tilapia

### Association between environmental variables and *An. funestus* larval abundance

The various environmental variables in the larval habitats associated with the presence of *An. funestus* larvae and other mosquito species are summarized in Table 2

**Table 2:**
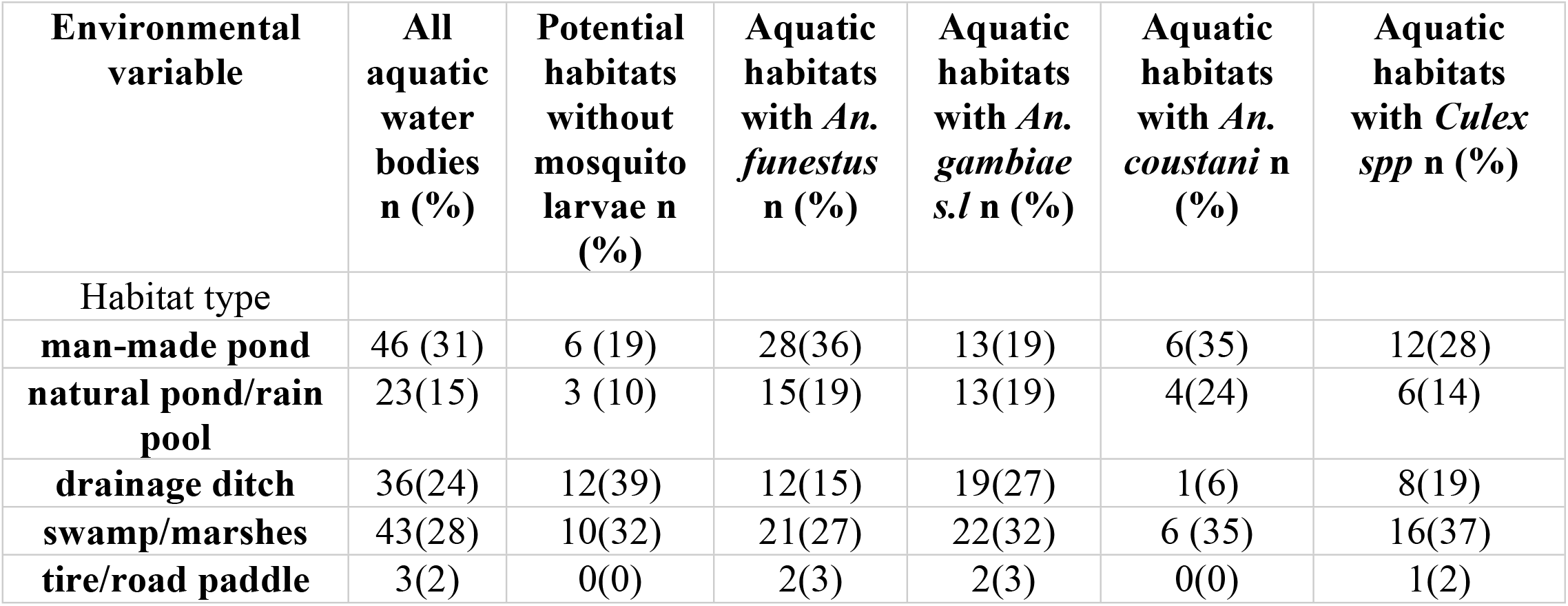
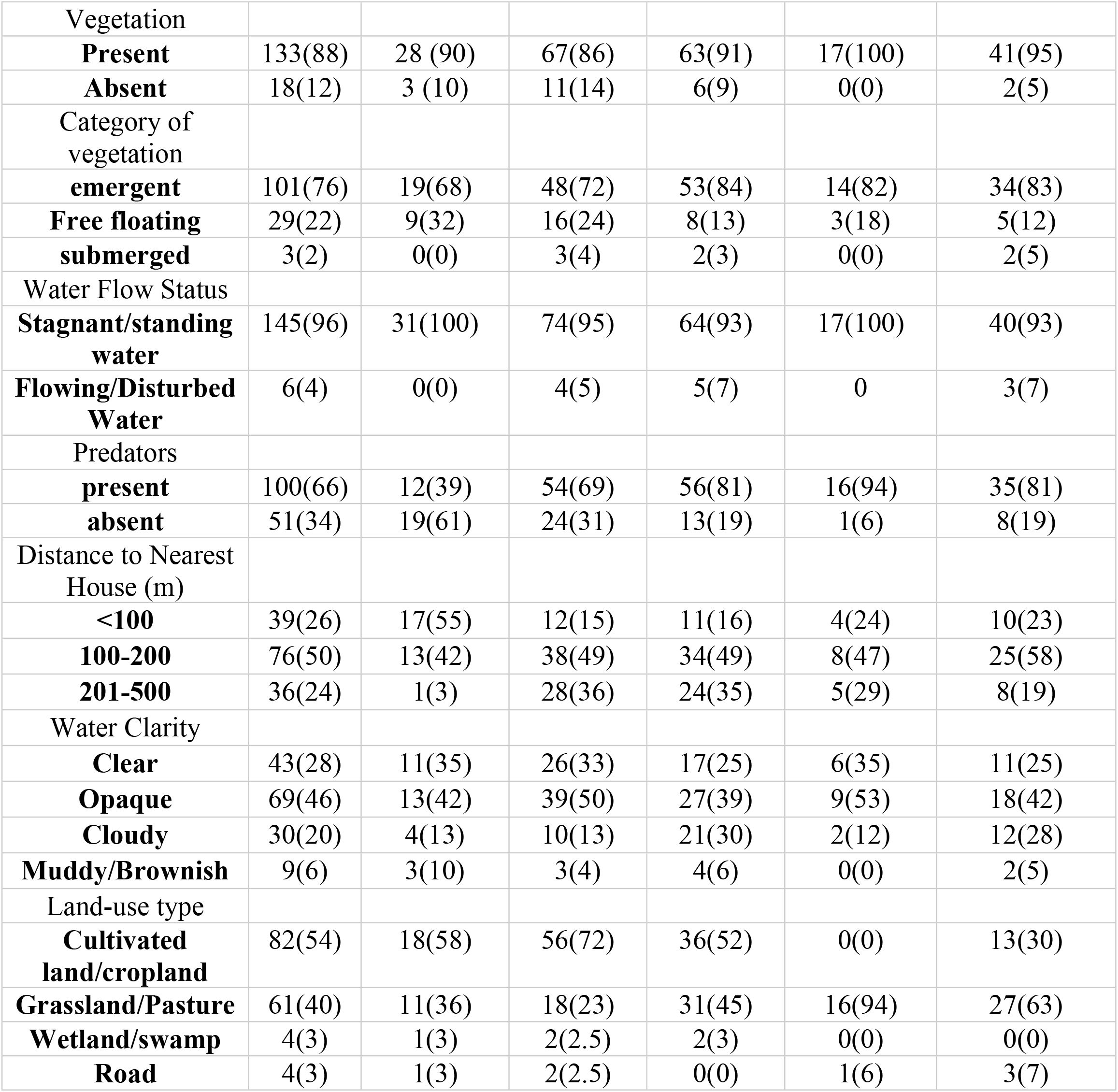
Characteristics of aquatic habitats of *An. funestus* and other mosquito species at Bungoma and Kombewa western Kenya (November 2019-November, 2020)

| Environmental variable | All aquatic water bodies n (%) | Potential habitats without mosquito larvae n (%) | Aquatic habitats with *An. funestus* n (%) | Aquatic habitats with *An. gambiae s.l* n (%) | Aquatic habitats with *An. coustani* n (%) | Aquatic habitats with *Culex spp* n (%) |
| --- | --- | --- | --- | --- | --- | --- |
| Habitat type |  |  |  |  |  |  |
| **man-made pond** | 46 (31) | 6 (19) | 28(36) | 13(19) | 6(35) | 12(28) |
| **natural pond/rain pool** | 23(15) | 3 (10) | 15(19) | 13(19) | 4(24) | 6(14) |
| **drainage ditch** | 36(24) | 12(39) | 12(15) | 19(27) | 1(6) | 8(19) |
| **swamp/marshes** | 43(28) | 10(32) | 21(27) | 22(32) | 6 (35) | 16(37) |
| **tire/road paddle** | 3(2) | 0(0) | 2(3) | 2(3) | 0(0) | 1(2) |
| Vegetation |  |  |  |  |  |  |
| **Present** | 133(88) | 28 (90) | 67(86) | 63(91) | 17(100) | 41(95) |
| **Absent** | 18(12) | 3 (10) | 11(14) | 6(9) | 0(0) | 2(5) |
| Category of vegetation |  |  |  |  |  |  |
| **emergent** | 101(76) | 19(68) | 48(72) | 53(84) | 14(82) | 34(83) |
| **Free floating** | 29(22) | 9(32) | 16(24) | 8(13) | 3(18) | 5(12) |
| **submerged** | 3(2) | 0(0) | 3(4) | 2(3) | 0(0) | 2(5) |
| Water Flow Status |  |  |  |  |  |  |
| **Stagnant/standing water** | 145(96) | 31(100) | 74(95) | 64(93) | 17(100) | 40(93) |
| **Flowing/Disturbed Water** | 6(4) | 0(0) | 4(5) | 5(7) | 0 | 3(7) |
| Predators |  |  |  |  |  |  |
| **present** | 100(66) | 12(39) | 54(69) | 56(81) | 16(94) | 35(81) |
| **absent** | 51(34) | 19(61) | 24(31) | 13(19) | 1(6) | 8(19) |
| Distance to Nearest House (m) |  |  |  |  |  |  |
| **<100** | 39(26) | 17(55) | 12(15) | 11(16) | 4(24) | 10(23) |
| **100-200** | 76(50) | 13(42) | 38(49) | 34(49) | 8(47) | 25(58) |
| **201-500** | 36(24) | 1(3) | 28(36) | 24(35) | 5(29) | 8(19) |
| Water Clarity |  |  |  |  |  |  |
| **Clear** | 43(28) | 11(35) | 26(33) | 17(25) | 6(35) | 11(25) |
| **Opaque** | 69(46) | 13(42) | 39(50) | 27(39) | 9(53) | 18(42) |
| **Cloudy** | 30(20) | 4(13) | 10(13) | 21(30) | 2(12) | 12(28) |
| **Muddy/Brownish** | 9(6) | 3(10) | 3(4) | 4(6) | 0(0) | 2(5) |
| Land-use type |  |  |  |  |  |  |
| **Cultivated land/cropland** | 82(54) | 18(58) | 56(72) | 36(52) | 0(0) | 13(30) |
| **Grassland/Pasture** | 61(40) | 11(36) | 18(23) | 31(45) | 16(94) | 27(63) |
| **Wetland/swamp** | 4(3) | 1(3) | 2(2.5) | 2(3) | 0(0) | 0(0) |
| **Road** | 4(3) | 1(3) | 2(2.5) | 0(0) | 1(6) | 3(7) |

Multiple regression analysis models revealed that distance to the nearest house (*P*=0.0122) was the best predictor of *An. funestus* larval density in the habitats (Table 3). The F-ratio in the ANOVA table revealed that, the *An. funestus* larval density was significantly associated statistically with key habitat variables (habitat type, land use type, vegetation coverage, vegetation height, distance to house, habitat size, water depth, water clarity, algae abundance and predator counts) at the time of larval sampling (*F*=2.06, d.f=10, 114, *P*=0.033, R^2^= 0.153). Furthermore, principal component analysis identified the distance to the nearest house as the key environmental factor associated with the abundance of *An. funestus* larvae.

**Table 3:** Multiple Regression analysis of factors associated with *An. funestus* larval density

| Habitat variable | Beta | Std.Err. of Beta | B | Std.Err. of B | t(114) | P-level |
| --- | --- | --- | --- | --- | --- | --- |
| Habitat Type | -0.0876 | 0.0984 | -0.0613 | 0.0688 | -0.8902 | 0.3752 |
| Land use type | -0.1816 | 0.1047 | -0.2464 | 0.1420 | -1.7348 | 0.0855 |
| Vegetation coverage | 0.0957 | 0.0997 | 0.0024 | 0.0025 | 0.9598 | 0.3392 |
| **Vegetation Height** | -0.0219 | 0.0890 | -0.0058 | 0.0236 | -0.2466 | 0.8056 |
| **Distance to house** | 0.2355 | 0.0925 | 0.0016 | 0.0006 | 2.5468 | *0.0122 |
| **Habitat size** | -0.0594 | 0.0929 | -0.0006 | 0.0009 | -0.6391 | 0.5240 |
| **Water depth** | 0.1088 | 0.0894 | 0.2402 | 0.1975 | 1.2163 | 0.2264 |
| **Water clarity** | 0.0171 | 0.1065 | 0.0033 | 0.0207 | 0.1605 | 0.8728 |
| **Algae abundance** | -0.1588 | 0.0980 | -0.0014 | 0.0008 | -1.6193 | 0.1081 |
| **Predator counts** | -0.0568 | 0.0909 | -0.0045 | 0.0072 | -0.6254 | 0.5329 |

Beta standardized coefficient, B the unstandardized coefficient value, \* significant at $P < 0.05$

The Mann–Whitney U and Kruskal–Wallis tests showed that *An. funestus* larval density had no significant difference between present and absent of vegetation (U=1192, *P*=0.978) (Fig. 4A), emergent and non-emergent vegetation (U=1369, *P*=0.166) (Fig. 4A), stagnant and flowing water (U=287, *P*=0.136) (Fig 5C), present and absent of aquatic predators (U=2215, *P*=0.162) (Fig 5B), among clear, opaque, cloudy and muddy/brownish water (χ2=7.316, df=3, *P*=0.062) (Fig 4C), or with the presence of *Pennisetum purpureum, Schoenoplectus californiscus* and other aquatic plant species in the habitats (χ2=2.671, df=2, *P*= 0.263) (Fig 4B) respectively. However, *An. funestus* larval density showed statistically significant differences associated with distances to the nearest house (<100, 100-200 and 201-500 m) (χ2=25.138, df=2, *P*<0.0001) (Fig. 5A) and land-use types (cultivated land/cropland, grassland/pasture, wetland/swamp and road) (χ2=29.197, df=3, *P*=0.000) (Fig 4C). Our field observation showed that *An. funestus is* most abundant in habitats surrounded by cultivated land compared to those in the grassland areas, wetlands, and roads.

**Fig 4:**
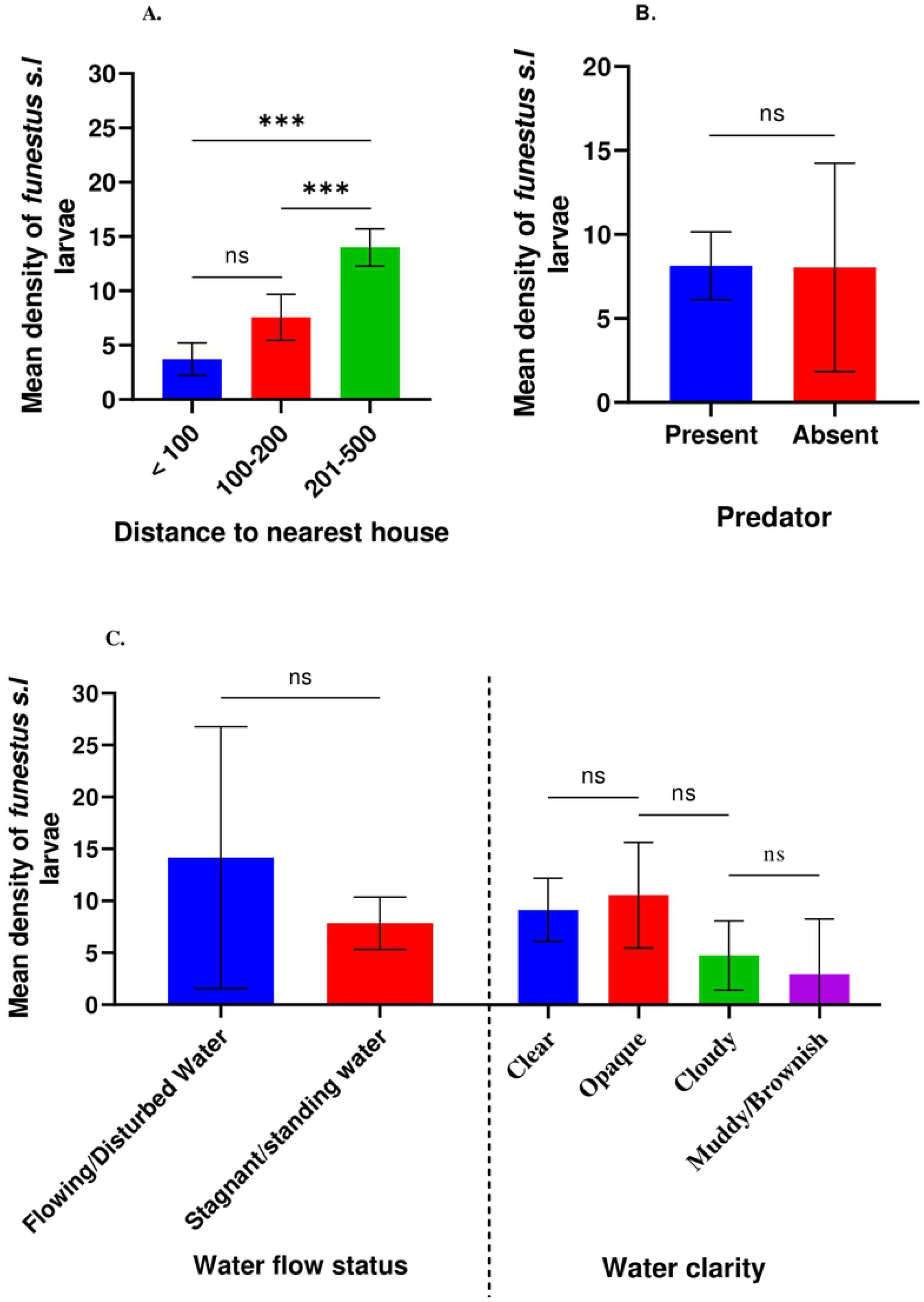
Association of habitat variables with *An. funestus* density (A) vegetation (present and absent) and category of vegetation (emergent and non-emergent), (B) aquatic species in the habitats, (C) land use type

**Fig 5:**
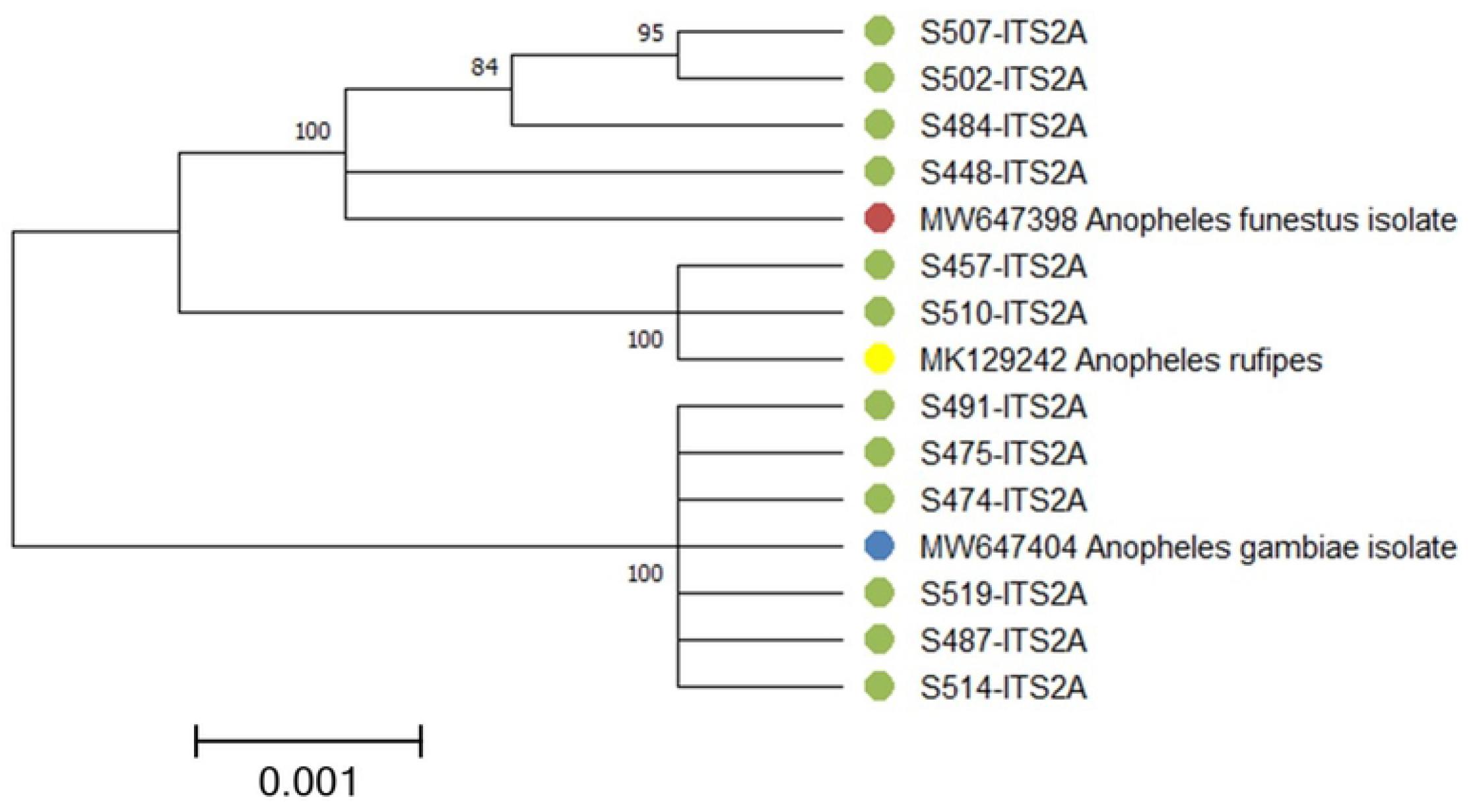
Association of habitat variables with *An. funestus* larval density (A) distance to the nearest house, (B) predator (present and absent in habitats) and (C) water flow status and water clarity

### Relationship between larval abundance and adults mosquitoes sampled indoors

The results of the Pearson correlation test showed that there was a statistically significant but weakly positive linear relationship between *An. funestus* larval abundance and the presence of adult mosquitoes collected indoors in Bungoma (rs=0.178, *P*=0.026). However, there was no significant relationship between *An. funestus* larval abundance and adult mosquitoes sampled indoors in Kombewa (rs=0.003, *P*=0.972).

### Composition of indoor resting *Anopheles* mosquitoes

A total of 1,221 *An. funestus s*.*l* and 195 *An. gambiae s*.*l* were collected indoors in Bungoma and Kombewa. Of the 1,221 *An. funestus s*.*l* sampled, 46% (n=565) and 54% (n=656) were collected from Bungoma and Kombewa respectively. For molecular identification, a sub-sample of 551 *Anopheles* comprising 380 *An. funestus s*.*l* and 171 *An. gambiae s*.*l* from the study sites were analysed. Of the *An. funestus s*.*l*. analysed, 201 and 179 were from Bungoma and Kombewa, respectively. Our results revealed that *An. funestus* predominated in the two study areas: *An. funestus* constituted 87% (n=201) and 88% (n=179) in Bungoma and Kombewa, respectively, while *An. rivulorum* constituted 13% (n=201) and 12% (n=179) for Bungoma and Kombewa, respectively.

Of the 171 *An. gambiae* s.l. analysed, 76 and 95 were from Bungoma and Kombewa, respectively. *An. gambiae* was the main species identified in Bungoma (86%, n= 76) and Kombewa (81%, n=95). *An. arabiensis* made up 15% (n=76) and 19% (n=95) in Bungoma and Kombewa, respectively. The evolutionary relationship of the unamplified sequenced data is shown in Fig 6.

**Fig 6:**
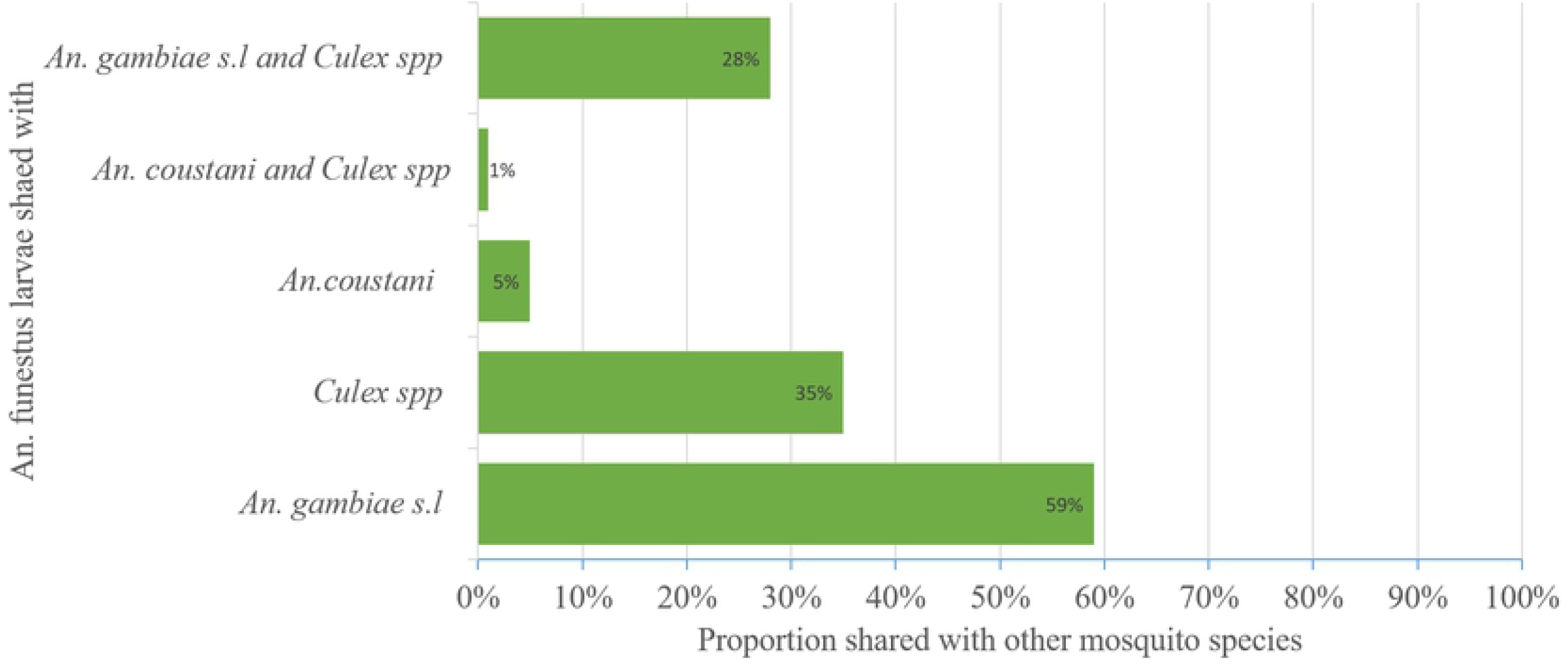
Evolutionary relationships of taxa.

The evolutionary history was inferred using the Neighbor-Joining method [37]. The optimal tree with the sum of branch length = 3.05389016 is shown (Fig 6). The percentage of replicate trees in which the associated taxa clustered together in the bootstrap test (1000 replicates) are shown next to the branches [38]. The evolutionary distances were computed using the Kimura 2-parameter method and are in the units of the number of base substitutions per site. The rate variation among sites was modelled with a gamma distribution (shape parameter = 1). This analysis involved 15 nucleotide sequences. All ambiguous positions were removed for each sequence pair (pairwise deletion option). There was a total of 815 positions in the final dataset. Evolutionary analyses were conducted in MEGA X [39].

### *Plasmodium falciparum (Pf)* sporozoite infection rate

All the sub-samples (n=551) subjected to molecular identification were genotyped to detect sporozoite. The *Pf* sporozoite rate of *An. funestus* in Bungoma and Kombewa was 2% (3/174) and 1% (2/157), respectively (Table 4). However, none of the *An. rivulorum* found in Bungoma and Kombewa was positive for *Pf* sporozoite (Table 4). Only one *An. gambiae* mosquito from Kombewa (1%, 1/77) tested positive for *Pf* sporozoite. However, sporozoites were not detected in *An. arabiensis* in the study sites.

**Table 4:** Sporozoite rate of *Anopheles* mosquitoes sampled from indoors in Bungoma and Kombewa, November 2019 to November 2020

| Site | *An. gambiae s.l* | No. tested | *Pf*+ve (%) | *An. funestus s.l* | No. tested | *Pf*+ve (%) |
| --- | --- | --- | --- | --- | --- | --- |
| Bungoma | *An. gambiae* | 65 | 0 (0) | *An. funestus* | 174 | 3 (2) |
|  | *An. arabiensis* | 11 | 0 (0) | *An. rivulorum* | 27 | 0 (0) |
|  | **Total** | **76** |  |  | **201** |  |
| Kombewa | *An. gambiae* | 77 | 1 (1) | *An. funestus* | 157 | 2 (1) |
|  | *An. arabiensis* | 18 | 0 (0) | *An. rivulorum* | 22 | 0 (0) |
|  | **Total** | **95** |  |  | **179** |  |

Numbers in the bracket indicate sporozoite rate; *Pf*, *Plasmodium falciparum*, +ve, positive

### *Anopheles* blood meal origins

A total of 208 samples (Bungoma, n=114 and Kombewa, n=94) were analysed for the origin of the mosquito blood meals. The HBI of *An. funestsus* was 84% (48/57) and 89% (39/44) for Bungoma and Kombewa, respectively (Table 5). Table 5 shows the HBI for all species tested.

**Table 5:** Blood meal sources of *Anopheles* mosquitoes sampled from indoors in Bungoma and Kombewa, November 2019 to November 2020

| Site | Species | No. tested | Blood meal sources |  |  | HBI |
| --- | --- | --- | --- | --- | --- | --- |
|  |  |  | human | bovine | goat |  |
| **Bungoma** | *An. gambiae* | 43 | 21 | 15 | 7 | 49 |
|  | *An. arabiensis* | 6 | 4 | 2 | 0 | 67 |
|  | *An. rivulorum* | 8 | 6 | 2 | 0 | 75 |
|  | *An. funestus* | 57 | 48 | 7 | 2 | 84 |
|  | **Total** | **114** | **79** | **26** | **9** |  |
| **Kombewa** | *An. gambiae* | 34 | 20 | 10 | 4 | 59 |
|  | *An. arabiensis* | 3 | 3 | 0 | 0 | 100 |
|  | *An. rivulorum* | 13 | 10 | 3 | 0 | 77 |
|  | *An. funestus* | 44 | 39 | 3 | 2 | 89 |
|  | **Total** | **94** | **72** | **16** | **6** |  |

HBI, human blood index

## Discussion

The great diversities of anopheline larval habitats in addition to their inaccessibility makes larval ecology studies of malaria mosquitoes methodologically cumbersome [15]. The presence of quality larval habitats is significant in determining the abundance and distribution of adult mosquitoes. This study was designed to add to the limited amount of information on the larval ecology of *An. funestus* in western Kenya. Two study areas were selected, a highland site (Bungoma) and a lowland site (Kombewa), and their aquatic habitats were examined and characterized to determine if there have been changes in the breeding habitats of *An. funestus* in the village sites. This study revealed that *An. funestus* is a major vector influencing malaria transmission in the region, confirming a previous report that *An. funestus* has re-emerged and could be responsible for malaria transmission in western Kenya [12].

Our findings revealed that *An. funestus* larvae thrive in a wide range of aquatic habitats and co-breeds with other malaria vectors in the same habitats. Although there were no significant differences observed in the various habitats types, man-made ponds had the highest proportion of *An. funestus* larvae. Man-made ponds, created mostly for making clay pots and sand winning, contributed remarkably to the proportion of *An. funestus* habitats and larvae abundance in the study areas. This corroborates previous findings in western Kenya where man-made habitats accounted for an increase in populations of *An. gambiae* [40,41]. Field observations have shown that man-made ponds are permanent habitats that hold water for a longer period compared to other habitat types. This suggests that malaria transmission in the study areas is partly man-made, and thus, proper environmental management specifically through habitat manipulation could curtail malaria transmission by major vectors of human malaria. This study confirms how anthropogenic modification of ecosystems can contribute greatly to the abundance and distribution of malaria vectors [42].

We found that more than 50% of *An. funestus* larvae co-existed in aquatic habitats with *An. gambiae s*.*l* larvae. Moreover, *An. funestus* shared the same habitats with *Culex spp* and *An. coustani*. Previous studies in neighbouring countries, Tanzania [43] and Uganda [44] have reported that *An. funestus* shared habitats with other *Anopheles* and *Culex spp*., indicating that any larval control programme targeting *An. funestus* would have a profound effect in controlling other equally important vectors of human malaria and other mosquito-borne diseases.

Hitherto, it has been reported that *An. funestus* prefers breeding in aquatic habitats with thick vegetation [15,16] but this study revealed that *An. funestus* can breed in habitats with aquatic vegetation or without vegetation. The presence of vegetation has been noted to be an important environmental variable associated with *Anopheles* mosquito larvae density [45]. For example, aquatic macrophytes play important role in the oviposition, larval survival and development of anophelines as they serve as a food source, protection for the larval stages and provide enabling environment for mosquito breeding [46–49]. However, this study revealed that *An. funestus* can breed in habitats with or without aquatic vegetation. Our data showed that there was no significant difference in the means between habitats with aquatic vegetation and habitats without aquatic vegetation.

Multiple linear regression analysis showed that the abundance of algae in the habitat was not a predicting factor for the density of *An. funestus* in this study. However, Gimnig et al [15] noted that algae abundance was positively correlated with *An. gambiae* density in western Kenya and also an important factor predicting the abundance of *An. pseudopunctipennis* in the Americas [50] and crucial for the development of *An. pretoriensis* in the Rift Valley province of Ethiopia [51].

While Aklilu et al [51] noted that algae abundance was an important factor, distance to the nearest house was not an important component associated with *An. gambiae s*.*l* and *An. pretoriensis* abundance in their study. However, the proximity of productive larval habitats to human or animal habitation to obtain a blood meal can determine the density of adult mosquitoes [52]. In western Kenya, a previous study reported that the distance to the nearest house was significantly associated with the abundance of *An. gambiae* [53]. Among the environmental variables assessed in our study, principal component and multiple linear regression analyses identified the distance to the nearest house as a major predictor of *An. funestus* abundance in habitats, in agreement with Minakawa et al [53] for *An gambiae*. Our findings suggest that larval source management targeting *An. funestus* aquatic habitats located near houses could reduce the adult mosquito population.

Aquatic predators are well known to influence the abundance of mosquito larvae in breeding environments and are considered beneficial biological control agents of mosquito larvae [54–57]. Notwithstanding, there was no significant difference in *An. funestus* larval density between aquatic habitats with predators and habitats without predators. A similar study by Ndenga et al [58] noted that the presence of predators was not significantly associated with the low density of *An. Gambiae s*.*l* larvae. Conversely, a previous study in Tanzania [59] and central Sudan [60] reported that most predators were identified in habitats with fewer densities of mosquito larvae. However, we witnessed that there was no reduction in the density of *An. funestus* larvae in the presence of predators in shared habitats. This could be ascribed to the presence of other prey in larval habitats. Kumar *et al* [61] documented that in the presence of alternative prey, the habitats, the larval consumption-ability of predators was significantly reduced in the habitats.

This study revealed that *An. funestus* was the predominant indoor resting vector corroborating the findings of previous investigations in Bungoma [19,20] and Kombewa [12,22]. The relative abundance, high sporozoite rate and HBI of *An. funestus* suggest that it is the main vector mediating malaria transmission in the study areas. We speculate that the adaptation of *An. funestus* to breed in warmer, open sunlit habitats may significantly reduce the developmental time of larval stages and increase the adult population of this species.

Because of a recent increase of 1° C due to climate change in Bungoma, its temperatures at 1,500 m asl are now equivalent to those at 1,347 m asl, assuming an altitudinal lapse rate of 154 m per degree Celsius at this latitude [62]. Thus, ecologically, an elevation of 1,500 m asl must now be treated as equivalent to a Kombewa’s low altitude vector ecosystem in western Kenya [63,64]. Essentially, climate change has made the malaria risk for the Kenya highlands similar to lowland habitats.

We acknowledge, however, the following limitations of our study: first, this study did not integrate detailed water chemistry analysis as part of the variable in assessing the larval ecology of *An. funestus*. Hence, further studies should be undertaken to assess the physicochemical characteristics of aquatic habitats that allow the co-existence of *An. funestus* with other mosquito species. Second, we were unable to examine the productivity of *An. funestus* aquatic habitats and their ability to facilitate the development of larvae to emerged adults.

## Conclusion

*An. funestus* was found breeding in a variety of aquatic habitats and co-existing with larvae of other mosquito species and aquatic predators. *An. funestus* was found in permanent and semi-permanent aquatic habitats, with or without aquatic vegetation, slow-moving/disturbed or stagnant water that was clear, opaque, cloudy and brownish. The only significant factor predicting the abundance *of An. funestus* in the aquatic habitats was the distance to the nearest house. Thus, larval control programmes should aim at targeting aquatic habitats near human dwellings to reduce the abundance of adult *An. funestus*. This study serves as a guide for the control of aquatic stages of *An. funestus* using larval source management or larviciding to complement existing vector control strategies. The relative abundance, high sporozoite rate, and HBI also confirm the importance of *An. funestus* in malaria transmission and the need for continuous vector surveillance before implementing vector control interventions.

## Acknowledgement

We extend our sincerest gratitude to all community leaders, farmland owners and heads of households of the Bungoma and Kombewa communities for permitting us to collect mosquitoes and larvae from their houses and farmlands.

